# Efficient biosynthesis of 3-hydroxypropionic acid in recombinant *Escherichia coli* by metabolic engineering

**DOI:** 10.1101/2025.08.26.672340

**Authors:** Yanwei Wang, Chenmei Suo, Jingyu Yang, Youxin Cui, Mutaz Mohammed Abdallah, Hongyi Yang, Pengchao Wang, Lixin Li, Changli Liu

**Author notes:** Correspondence to: Changli Liu, *E-mail* addresses, Lixin Li, *E-mail* addresses. These authors contributed equally (Yanwei Wang, Chenmei Suo, Jingyu Yang, Youxin Cui). Key Laboratory of Saline-alkali Vegetation Ecology Restoration, Ministry of Education, College of Life Sciences, Northeast Forestry University, Harbin 150040, China.

## Abstract

3-Hydroxypropionic acid (3-HP) is a platform compound that can produce many chemical commodities. This study focuses on establishing and optimizing the production of 3-HP in *E. coli*. We constructed a series of engineered *E. coli* strains which can produce 3-HP via the malonyl-CoA pathway. To increase the metabolic flux of acetyl-CoA, a precursor for the synthesis of 3-HP, CRISPR/Cas9-based DNA editing technique was used to knock out the genes encoding pyruvate oxidase (*poxB)*, lactate dehydrogenase (*ldhA*) and phosphate transacetylase (*pta*), thereby reducing the formation of by-products. Concurrently, the acetyl coenzyme a carboxylase gene (*accDABC*) is overexpressed on the chromosome with the objective of augmenting intracellular acetyl-CoA levels and, consequently, 3-HP production. Next, we introduced a plasmid containing a codon-optimized malonyl-CoA reductase gene (*mcr*) into the engineered strain. Finally, we constructed a transcription factor-based metabolite biosensor utilizing the PpHpdR/P*hpdH* system, followed by the screening of mutant strains for enhanced 3-HP production through adaptive laboratory evolution. Combining the above metabolic engineering efforts with optimisation of media and fermentation conditions, the 3-HP titer of the engineered strain WY7 increased from an initial titer 0.34 g/L to 48.8 g/L. This study encourages further research in metabolic pathway optimizationto produce 3-HP.

**Highlights:**

- Synthesis 3-HP in the malonyl-CoA pathway.
- Edit the *Escherichia coli* genome using the CRISPR/Cas9 system.
- Elevated production of 3-HP by knocking out bypass genes *ldhA*/*pta*/*poxB*.
- A biosensor was designed to respond to 3-HP concentration.
- Adaptive laboratory evolutionary strategies increase 3-HP production.

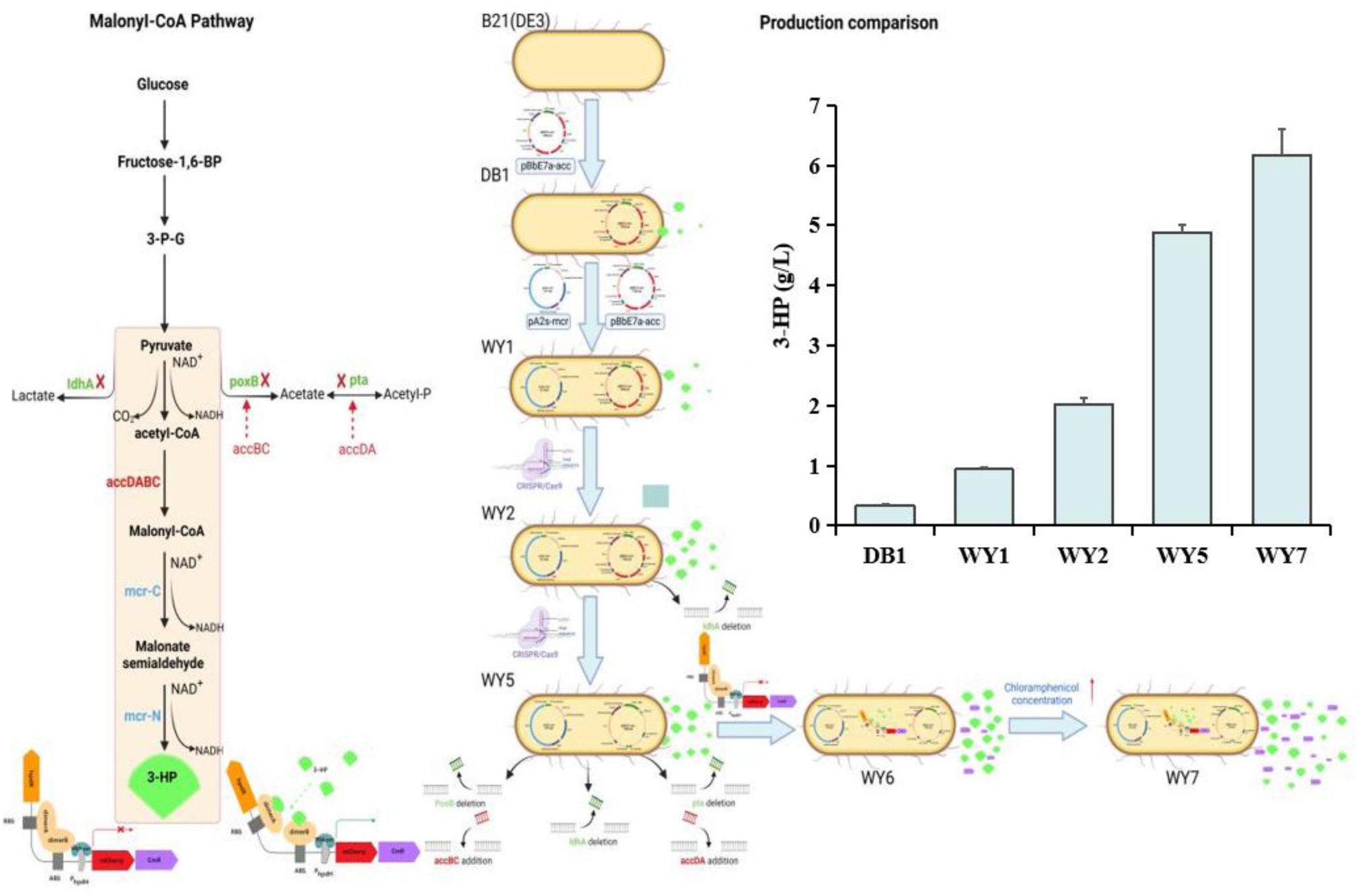

## 1. Introduction

3-Hydroxypropionic acid (3-HP), chemically represented as C_3_H_6_O_3_, is also referred to as β-hydroxy propionic acid. This compound is a clear, viscous liquid that lacks any noticeable odor and blends seamlessly with water, alcohol, ether, and various organic solvents [1]. Its versatility allows it to serve as a food and feed additive or preservative [2]. Additionally, 3-HP exhibits targeted nematocidal properties, particularly against Meloidogyne incognita, a plant-parasitic nematode, making it suitable for formulating nematicides [3].

3-HP exhibits strong chemical reactivity owing to the presence of both hydroxyl and carboxyl functional groups. While it shares isomerism with lactate, 3-HP demonstrates greater reactivity due to the distinct positioning of its hydroxyl group [4]. Furthermore, 3-HP serves as a versatile precursor in chemical synthesis, enabling the production of compounds such as 1,3-propanediol (1,3-PDO), 3-hydroxypropionaldehyde (3-HPA), acrylic acid, and malonic acid^[5^, ^6, 7]^. It also acts as a key substrate for cyclization and polymerization processes, yielding products like propiolactone, polyesters, and oligomers. Through polymerization of its hydroxyl and carboxyl groups, 3-HP can form poly(3-hydroxypropionate) (Poly(3HP))^[8]^. Poly(3HP) is a well-studied biodegradable polyester known for its biocompatibility and mechanical strength [9]. These properties make it suitable for medical applications, including the fabrication of prosthetics and heart valves, highlighting its potential in the healthcare sector [10]. Due to its advantageous characteristics, 3-HP was recognized in a 2004 U.S. Department of Energy report as one of the 12 most promising chemical compounds globally [11].

Initially, 3-hydroxypropionic acids are produced through both chemical and biological methods. However, chemical synthesis is often associated with environmental concerns such as high energy demands, complex purification steps, and the generation of persistent waste by-products [12]. In contrast, biological production offers a more sustainable alternative, minimizing ecological impact. As a result, biosynthetic approaches to 3-HP production have gained attention as a foundation for environmentally friendly, large-scale manufacturing.

The early discovery of 3-HP biosynthesis was reported by Sobolov et al. [13], who observed that a wild-type *Lactobacillus* strain could generate 3-HP when grown on glycerol. This finding laid the foundation for subsequent research into microbial 3-HP production. A widely adopted approach for scaling up industrial 3-HP synthesis involves the genetic modification of metabolic pathways [14]. Engineered strains, optimized through these techniques, demonstrate enhanced efficiency in converting low-cost carbon substrates into 3-HP compared to their wild-type counterparts [15]. Currently, the primary biosynthetic routes for 3-HP include the glycerol, β-alanine, and malonyl-CoA pathways [16].

The malonyl-CoA pathway offers distinct advantages over alternative routes for 3-HP production. Unlike other pathways, it eliminates the requirement for costly coenzyme B12 supplementation and utilizes economical carbon substrates [17]. Initially, acetyl-CoA is transformed into malonyl-CoA by acetyl-CoA carboxylase (ACC), a multimeric enzyme composed of ACCA, ACCB, ACCC, and ACCD subunits [4]. Malonyl-CoA is then reduced to 3-HP by malonyl-CoA reductase (MCR). Due to its efficiency and reduced production costs, this route has attracted widespread interest for sustainable 3-HP synthesis [18]. Furthermore, ACC functions as a crucial rate-limiting enzyme, playing a significant role in regulating 3-HP accumulation through this pathway [4].

The overexpression of acc can enhance malonyl-CoA accumulation in cells [19], hence augmenting the generation of the downstream product. Consequently, the overexpression of the endogenous gene acc from *E. coli*, along with *mcr* from *Chloroflexus aurantiacus* in *E. coli* K12, resulted in the effective construction of an engineered strain with a 3-HP titer of 0.128 g/L, which was 2-fold higher compared to the original strain without overexpression of *acc* [15]. However, overexpression of *acc* producing excessive accumulation of malonyl-CoA will inhibit the growth of cells [20]. Wang et al. enhanced 3-HP production and cell growth by modulating the expression levels of the *acc* subunits (*accBC* and *DtsR1*), thereby alleviating malonyl-CoA-associated cytotoxicity. This optimization enabled *C. glutamicum* to achieve a 3-HP concentration of 6.8 g/L in shake-flask fermentation. To address the functional imbalance between the N- and C-terminal domains of malonyl-CoA reductase (MCR), Liu et al. divided the *mcr* gene into two separate coding regions [21]. By rebalancing the enzymatic activities of the MCR-C and MCR-N fragments, they achieved a dramatic 270-fold increase in 3-HP production, with final titers reaching 40.6 g/L [21].

Although experimental designs based on traditional metabolic engineering can meet most requirements, The intricacy of carbon and energy metabolism in microbial cells presents obstacles in regulating the titer of target products generated by microbes in certain instances[22]. The Adaptive Laboratory Evolution (ALE) technique can intentionally replicate the variation and selection processes of natural evolution in a laboratory setting [23]. ALE directs the evolutionary trajectory of microorganisms by applying artificial selection pressure, resulting in target strains with advantageous mutations through screening [24]. In contrast to conventional metabolic engineering, which considers the intricate metabolic network within the cell, the ALE technique merely requires the construction of relevant interference factors based on specific requirements [25]. The ALE approach facilitates the acquisition of high-titer strains of specified products through screening, presenting a wider scope for development [26].

ALE facilitated by metabolite biosensors utilizing transcription factors represent a highly effective approach for the efficient production of target metabolites [27, 28, 29]. In comparison to the conventional approach of adaptive laboratory evolution, the utilisation of biosensors can markedly enhance the screening throughput and selectivity, thereby expediting the evolution of strains.

In this study, the malonyl-CoA pathway for 3-HP biosynthesis was engineered in *E. coli* by co-expressing the native *accDABC* genes, encoding acetyl-CoA carboxylase, alongside a codon-optimized *mcr* gene from *C. aurantiacus*. To minimize the cellular burden associated with plasmid maintenance and enhance genetic stability, the CRISPR/Cas9 genome editing system was employed. Specifically, the genes encoding pyruvate oxidase (*poxB*), lactate dehydrogenase (*ldhA*), and phosphate acetyltransferase (*pta*) were disrupted, and the *accDABC* operon was chromosomally integrated to enable stable overexpression (Fig. 1). The resulting strain, designated Q1Z2, served as the host for transformation with a plasmid carrying *mcr*. Additionally, a 3-HP-responsive biosensor plasmid was developed, capable of linking 3-HP production to cell growth under chloramphenicol selection pressure. This biosensor enabled ALE, facilitating the enrichment of strains with enhanced 3-HP production capacity. Through iterative rounds of ALE and screening, we obtained a high-producing strain, WY7, which achieved a 3-HP concentration of 48.8 g/L in 5-liter fed-batch fermentation.

**Fig. 1.**
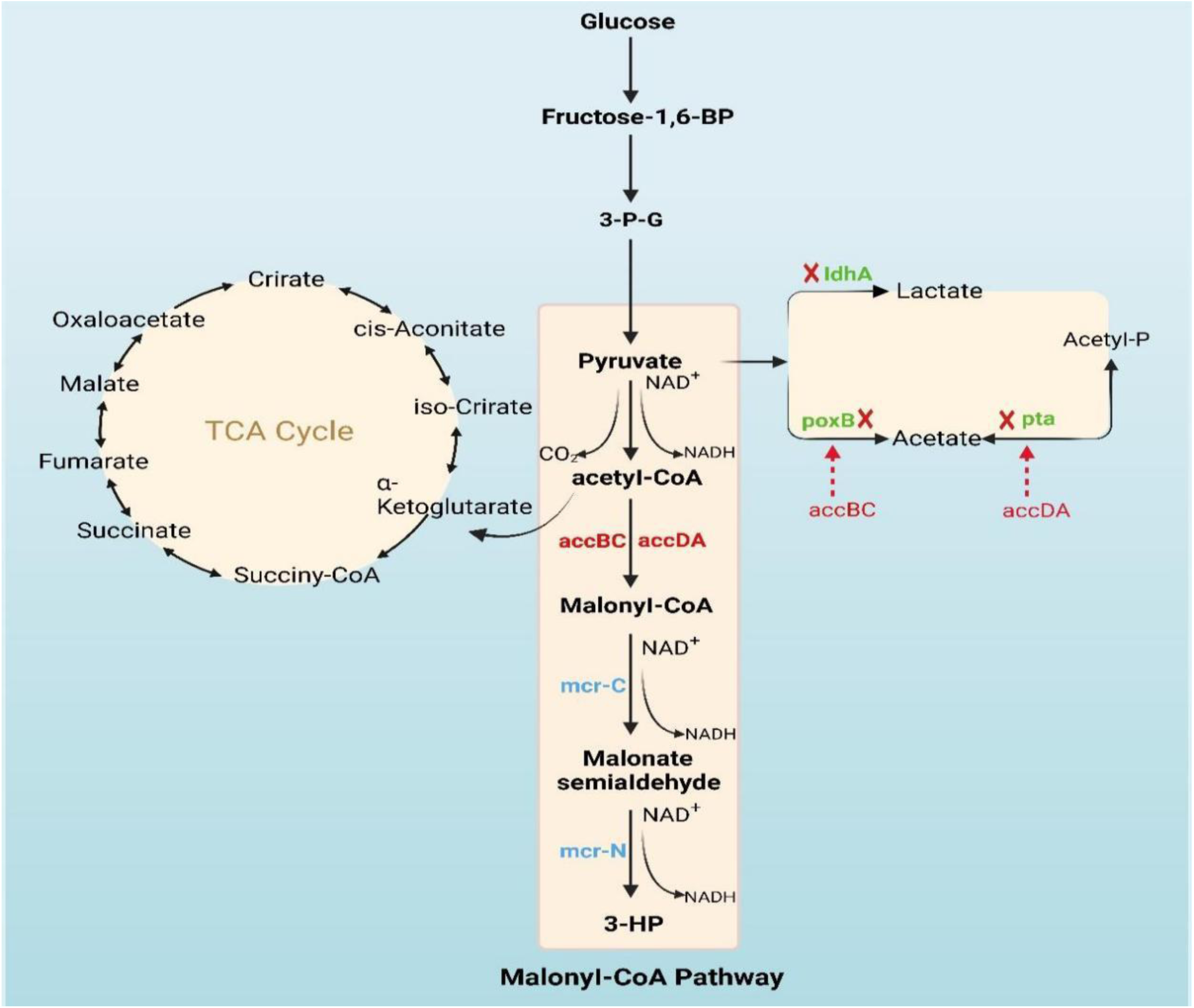
Engineering the malonyl-CoA pathway in *E. coli* for 3-HP biosynthesis. Endogenously overexpressed enzymes are indicated in red, heterologous enzymes in blue, and reactions associated with gene deletions in green. Red arrows denote integration of the corresponding gene at the site of the deleted gene. Enzyme abbreviations: *accDABC*, the four subunits (*accD, accA, accB, accC*) of acetyl-CoA carboxylase; *mcr-C,* C-terminal domain of the malonyl-CoA reductase; *mcr-N*, N-terminal domain of the malonyl-CoA reductase; *ldhA*, lactate dehydrogenase; *poxB*, pyruvate oxidase; *pta*, phosphate transacetylase.

## 2. MATERIAL AND METHODS

### 2.1 Bacterial strains, plasmids, and reagents

The bacterial strains and plasmids applied in this research are detailed in Table 1. *E. coli* DH5α was employed for the assembly and maintenance of all recombinant plasmids, whereas *E. coli* BL21(DE3) served as the expression host for protein production and 3-hydroxypropionic acid (3-HP) synthesis. Plasmid and genomic DNA extractions were carried out using the Plasmid Mini Kit and Bacterial DNA Kit (OMEGA), respectively. High-fidelity DNA amplification was performed using Phanta Flash Super-Fidelity DNA Polymerase, and Taq DNA Polymerase along with restriction enzymes were sourced from Vazyme (Nanjing, China) or New England Biolabs (Ipswich, USA). PCR-amplified DNA was purified using a Gel Extraction Kit (OMEGA). Gene fragment ligation with homologous ends was achieved using the ClonExpress Ultra One-Step Cloning Kit (Vazyme). The 3-HP standard compound was obtained from Sigma-Aldrich, and other reagents were supplied by Sangon Biotech (Shanghai, China).

**Tab. 1.** Experimental main plasmids and strains.

| plasmid/Strain | Relevant features/genotype | Reference |
| --- | --- | --- |
| **Plasmids** |  |  |
| pE7a-acc | ColE1 Amp P<sub>T7</sub>-accDABC | Preserve in lab |
| pA2c-rfp | P15A; PtetR; CmR | Preserve in lab |
| pE8c-mcr-PrpE | mcr; CmR | Preserve in lab |
| pLB1s | Str | Preserve in lab |
| pA2s-mcr | p15A Str P<sub>tet</sub>-mcr | This study |
| pA2s-mcr-N-C* | p15A Str P<sub>tet</sub>-mcr-N-mcr-C | This study |
| pTargetF | pMB1 aadA P<sub>J23119</sub>-sgRNA- | Preserve in lab |
| pTargetF-*ldhA* | pMB1 aadA P<sub>J23119</sub>-sgRNA-*ldhA* | This study |
| pTargetF-*poxB* | pMB1 aadA P<sub>J23119</sub>-sgRNA-*poxB* | This study |
| pTargetF-*pta* | pMB1 aadA P<sub>J23119</sub>-sgRNA-*pta* | This study |
| pCas | repA101(Ts)kan P<sub>cas</sub>-cas9 P<sub>araB</sub>-Red lacI<sup>q</sup> P<sub>trc</sub>-sgRNA-pMB1 | Preserve in lab |
| pYB-eGFP | p15A eGFP | Preserve in lab |
| pUAM_O63-mcherry | pUAM Amp mCherry | Preserve in lab |
| pYB-0055-mCherry | p15A Amp P<sub>HPDH</sub>-mCherry | This study |
| pYB-0055-mCherry-CmR | p15A Amp P<sub>HPDH</sub>-mCherry-CmR | This study |
| **Strains** |  |  |
| *E. coli* BL21(DE3) | F<sup>-</sup> *ompT hsdS<sub>B</sub>(r<sub>B</sub><sup>-</sup> m<sub>B</sub><sup>-</sup>) gal dcm rne131* (DE3) | Preserve in lab |
| *E. coli* DH5α | ΔM15, Δ(lacZYA -argF)U169, deoR, recA1, endA1, hsdR17 (rK<sup>-</sup>, mK<sup>+</sup>), phoA, supE44, λ<sup>-</sup>, thi<sup>-</sup>1, gyrA96, relA1 | Preserve in lab |
| BL21(DE3)/pCas | BL21(DE3) harboring plasmid pCas | This study |
| BW25113/pTargetF | BW25113 harboring plasmid pTargetF | Preserve in lab |
| BW25113/pCas | BW25113 harboring plasmid pCas | Preserve in lab |
| BL21Δ*ldhA* | BL21(DE3) Δ*ldhA* | This study |
| BL21Δ*poxB* | BL21(DE3) Δ*poxB* | This study |
| BL21Δ*pta* | BL21(DE3) Δ*pta* | This study |
| Q1Z1 | BL21(DE3) $\Delta ldhA \Delta pta::119-accDABC$ | This study |
| Q1Z2 | BL21(DE3) $\Delta ldhA \Delta pta::119-accDABC \Delta poxB::CPA1-accDABCaccBC$ | This study |
| DB1 | BL21(DE3)/ pA2s-mcr | This study |
| DB2 | BL21(DE3)/ pE7a-acc | This study |
| WY0 | DB1/pE7a-acc | This study |
| WY1 | DB2/pA2s-mcr-N-C* | This study |
| WY2 | WY1 with the deletion of *ldhA* | This study |
| WY3 | WY1 with the deletion of *poxB* | This study |
| WY4 | WY1 with the deletion of *pta* | This study |
| WY5 | Q1Z2/pA2s-mcr-N-C* | This study |
| CS1 | BW25113/pYB-0055-mCherry-CmR | This study |
| CS2 | Q1Z2/pYB-0055-mCherry-CmR | This study |
| WY6 | WY5/pYB-0055-mCherry-CmR | This study |
| WY7 | WY6 with ALE | This study |

### 2.2 SDS-PAGE for protein expression analysis

The expression of proteins encoded by the recombinant plasmids pE7a-acc and pA2s-mcr was evaluated in *E. coli* BL21(DE3). Cultures were incubated in LB medium at 37 °C, supplemented with appropriate antibiotics streptomycin (50 mg/L) and ampicillin (100 mg/L) and various concentrations of inducers (IPTG at 10, 100, and 200 μM; Tc at 10 nM). After a 14-hour induction period, cells were collected and lysed using ultrasonic disruption. The resulting lysates were centrifuged at 12,000 rpm for 10 minutes to separate soluble and insoluble fractions. Protein expression levels were assessed by 12% SDS-PAGE under denaturing conditions, followed by staining with Coomassie Brilliant Blue R-250 to visualize the protein bands.

### 2.3 Construction of plasmid expressing the *mcr*

The p15A replication origin, streptomycin resistance gene *Str*, Ptet+tetR promoter, and *mcr* gene were amplified with primers ZT-F/R, Str-F/R, Tet-F/R, and MCR-F/R using plasmids pA2c-*rfp*, pLB1s, and pE8c-mcr-prpE as templates, respectively, and the homology arms were ligated using the ClonExpress Ultra One Step Cloning Kit to construct the pA2s-mcr plasmid. The plasmid pA2s-mcr-N-C* was constructed using pA2s-mcr as a template. The primers MCR-C-F/MCR-N-R were employed to amplify the *mcr* gene, resulting in its fragmentation. Additionally, a ribosome binding site sequence was introduced at the 5’ end of the *mcr*-C fragment. Subsequently, mutations at positions N940V, K1106W, and S1114R were introduced into the segmented *mcr* plasmid using the primer pairs 940-F/1106-1114-R and 1106-1114-F/940-R.

### 2.4 Media and culture conditions for 3-HP production in the flask

All recombinant *E. coli* strains were cultured in LB medium at 37℃ overnight at 200 rpm. Then, 300 μL cultured medium was measured and transferred to 10 mL modified M9 medium (20 g/L glucose, 2 g/L yeast extract) in a 50 mL flask and grown at 37℃for 48 h at 200 rpm. 50 mg/L of streptomycin and 100 mg/L of ampicillin were added to the culture to select the plasmids. The recombinant cells were induced at 0.8-1.2 OD_600_ with 10 μM IPTG and 200 nM Tc, respectively. After 2.5 h of induction, 40 mg/L biotin and 20 mM NaHCO_3_ were added to the medium to assist the expression of *acc*. The antibiotics and inducer agents were supplied periodically after induction of IPTG every 12 h until 48 h.

### 2.5 Construction of the synthetic 3-HP selection vector

The plasmids pYB-eGFP and pUAM_O63-mcherry contained vector backbone and mCherry gene sequence were preserved in our laboratory, the synthesized plasmid pUC19-PP0055 contained the 3-HP response promoter sequence. These three plasmids were used as the template. The required vector and target fragment were amplified by three pairs of primers PYB-F/PYB-R, mCherry-F/mCherry-R, and 0055-F2/0055-R, which have 15-20 bp homologous sequences (see supplementary Tab. S1). The amplified bands were detected by electrophoresis, recovered and purified by Gel Extraction Kit (Omega, D2500-01). Clon Express Ultra One Step Cloning Kit (Vazyme, C113-01) was used to connect the purified fragments to obtain the recombinant plasmid pYB-0055-mCherry. Similarly, CmR was amplified and inserted downstream of mCherry to construct the final biosensor plasmid pYB-0055-mcherry-CmR.

Genome editing of the engineered *E. coli* was performed using a two-plasmid CRISPR/Cas9 system, following the method described by Jiang et al. [38]. This system utilizes two separate plasmids: pCas and pTargetF. The pCas plasmid harbors the *cas9* gene encoding the Cas9 endonuclease, a temperature-sensitive origin of replication, a kanamycin resistance marker, and the λ-Red recombination genes under the control of an arabinose-inducible promoter. Additionally, it includes a pMB1-sgRNA element regulated by IPTG for plasmid curing. The pTargetF plasmid carries the N20-sgRNA sequence specific to the target gene, along with a pMB1 origin of replication and a streptomycin resistance gene.

Specifically, we knocked out the *ldhA* genes of the *E. coli* BL21 (DE3) genome using CRISPR/Cas9 technology and verified whether the genes were successfully knocked out by colony PCR (see supplementary Fig. S2). Following the elimination of the plasmids pCas and pTargetF, which were employed in the knockout process. And then knocked out the *poxB* and *pta* genes, respectively. The pE7a-acc and pA2s-mcr-N-C* were transferred into these strains then we successfully constructed engineered strains WY2-WY4. Specifically, we expressed *accD* and *accA* using promoter P_119_ and integrated the gene expression unit into the original *pta* locus; we expressed *accB* and *accC* using promoter P_CPA1_ and incorporated it into the original *poxB* locus. Colony PCR and gene sequencing verified the gene editing sites of this strain. (see supplementary Fig. S3) Based on the above experiment, the 3-HP-producing strain WY5 was constructed by transferring the plasmid pA2s-mcr-N-C* into strain Q1Z2 with the genotype of *ΔldhAΔpta::accDABC ΔpoxB::accDABCaccBC*.

### 2.6 Adaptive Laboratory Evolution

A single colony of the WY6 strain was cultured in modified M9 medium (20 g/L glucose, 2 g/L yeast extract) supplemented with 50 mg/L streptomycin and 100 mg/L ampicillin. Upon reaching the stationary phase, the culture was diluted to an OD_600_ of 0.1 in fresh medium containing 200 nmol/L Tc and 136 mg/L chloramphenicol. The culture was subcultured every 12 h into fresh medium at an initial OD600 of 0.1, all experiments were performed in triplicate. With a constant Tc concentration, ALE was conducted on the original WY6 strain using a strategy that gradually increased chloramphenicol concentrations. After the final round, cells were plated on LB agar with antibiotics and incubated overnight at 37°C to isolate single colonies. The 3-HP titer in the fermentation broth was quantified using HPLC (High-performance liquid chromatography).

### 2.7 Fed-batch fermentation

The engineered strain was cultivated in a 5 L fermenter containing 2 L of modified minimal medium containing 20 g/L glucose, 2 g/L yeast extract, 9.8 g/L K_2_HPO_4_·3H_2_O, 3.0 g/L (NH_4_)_2_SO_4_, 2.1 g/L C_6_H_8_O_7_·H_2_O, 0.3 g/L ammonium ferric citrate, 0.5 g/L MgSO_4_, 9 mg/L CaCl_2_·2H_2_O, 6 mg/L FeSO_4_·7H_2_O, 2 mg/L H_3_BO_3_, 2 mg/L MnCl_2_·4H_2_O, 0.8 mg/L (NH_4_)_6_Mo_7_O_24_·4H_2_O, and 0.2 mg/L CuSO_4_·5H_2_O. The pH was maintained at 6.8 through automated addition of ammonia (28% v/v). Dissolved oxygen (DO) was kept at 25% by regulating agitation speed and airflow. Once the initial glucose was depleted, a concentrated glucose feed was introduced to sustain a residual concentration of 1–5 g/L. Induction of exogenous gene expression began at an OD600 of 12 using 200 nmol/L Tc, with additional Tc supplemented every 12 h.

### 2.8 Analytical methods

The cell density was measured in a 10-mm-path-length cuvette using a double-beam spectrophotometer (Tryte Technology, TR-TC-U4000). The concentrations of 3-HP in the culture broth were determined by HPLC (1525; Waters; CA; USA). Briefly, the supernatant obtained by centrifugation of the culture samples at 12,000 rpm for 10 mins was filtered using a 0.22 μm Nylon-membrane and eluted through a 300-mm × 7.8-mm Xtimate Sugar-H column. The mobile phase was 5 mmol/L H_2_SO_4,_ and the flow rate was 0.4 mL/min during elution. The column and cell temperatures were 65 and 45°C, respectively.

## 3. Results and Discussion

### 3.1 Construction of a malonyl-CoA route for 3-HP production in *E. coli*

Wild-type *E. coli* does not produce 3-HP, but synthesizes a small amount of malonyl-CoA, an important precursor of 3-HP, catalyzed by acetyl-CoA carboxylase. From the knowledge above, we postulated that by heterologously expressing the *mcr* gene in *E. coli*, it could synthesize 3-HP from glucose through the malonyl-CoA pathway. After overexpressing *accDABC*, the 3-HP production of the strain can be further increased. To construct the 3-HP synthesis pathway in *E. coli*, we constructed the recombinant plasmid pA2s-mcr according to the BioBricks™ standard proposed by Lee et al [30]. In this plasmid, we selected the promoter P_TetR_ to express the *mcr* gene. In addition, the streptomycin resistance gene *Str* and the medium-copy-number replication origin p15A were selected to form the plasmid backbone. P_TetR_ is an inducible promoter that, when tetracycline is present, can initiate the expression of downstream genes [31, 32]. Plasmid construction was tested by colony PCR, enzyme digestion validation (see supplementary Fig. S1) and sequencing. In contrast, plasmid pE7a-acc used the P_T7_ promoter induced by IPTG to express the *accDABC*, and its plasmid backbone consisted of ampicillin resistance gene Amp and high-copy-number ColE1 origin of replication. According to the BioBricks™ standard, these two plasmids can coexist. Then, the plasmids pA2s-mcr and pE7a-acc were transferred into *E. coli* BL21(DE3) respectively to construct engineering strain named DB1 and DB2. SDS-PAGE, shown in Fig. 2, detected these two strains’ protein expression.

**Fig. 2.**
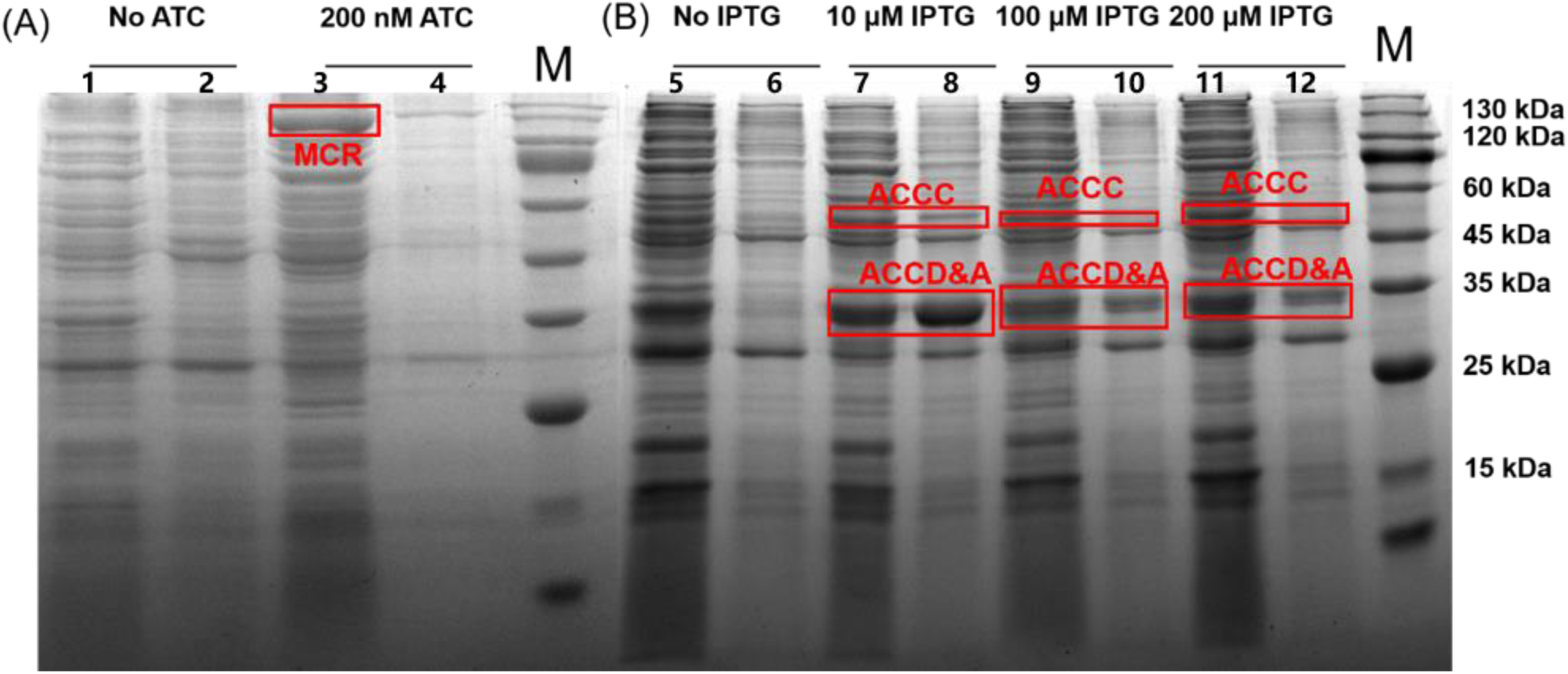
SDS-PAGE for protein expression analysis. (A) Expression of MCR protein (∼132.1 kDa) in recombinant *E. coli* DB1. M, protein marker. Lane1, the cell-free soluble extracts obtained from *E. coli* wild-type strain. Lane 2, the cell-free insoluble extracts obtained from *E. coli* wild-type strain. Lane 3, the cell-free soluble extracts obtained from DB1. Lane 4, the cell-free insoluble extracts obtained from DB1; (B) Effects of IPTG concentration on the expression of ACCDABC proteins in recombinant *E. coli* DB2. Lanes 5/7/9/11 were the cell-free soluble extracts; lanes 6/8/10/12 were the cell-free insoluble extracts. The upper arrow indicates ACCC (∼49.3 kDa), and the lower arrow represents ACCD (∼33.3 kDa) and ACCA (∼35.2 kDa); M, protein marker.

As shown in Fig. 2A, lane 3 shows the MCR protein band; MCR has an approximate molecular weight of 132.1 kDa, which band did not exist in the uninduced strain. Fig. 2B shows the expression of ACC, because of the approximate molecular weights are very similar to ACCA (35.2 kDa) and ACCD (33.3 kDa), these proteins were thought to be combined into a single band. Similarly, ACCC, the 49.3 kDa protein band, was seen on the protein band gel. The protein expressed by gene *accB* (16.2 kDa) was not shown obviously by bands, probably due to its low molecular weight in the front of the gel, which is consistent with the results obtained by Rathnasingh et al. (2012). After completing the SDS-PAGE assay, these two plasmids were co-transferred into *E. coli* BL21 (DE3) to construct strain WY0.

To verify the ability of the engineered strains of synthesizing 3-HP, we incubated strain WY0 and DB1 in 5 mL of LB liquid medium overnight at 37℃. Then, we moved the culture solution into 10 mL of modified M9 medium at 3% inoculum. After the strains were cultured in M9 medium at 37℃ for 48 hours, after the cells were collected, HPLC was used to detect the 3HP titer. The recombinant strains DB1 and WY0 could synthesize 3-HP of 0.15 g/L and 0.34 g/L, respectively. Compared with the recombinant strain DB1, the overexpression of the *acc* gene doubled the 3-HP titer of the engineered strain. The above results proved that we have successfully constructed the synthetic pathway of 3-HP in *E. coli*. To further increase the yield of 3-HP, the *mcr* gene was split into two segments [21] for separate expression. We then used the targeted mutation technique to introduce the mutation site (N940V/K1106W/ S1114R), which improves the activity of the MCR enzyme, to obtain the mutant plasmid pA2s-mcr-N-C*. The plasmid was transferred into the *E. coli* DB2, and the recombinant strain WY1 was obtained, which could synthesize 3-HP efficiently with a titer of 0.69 g/L (Fig. 3A).

**Fig. 3.**
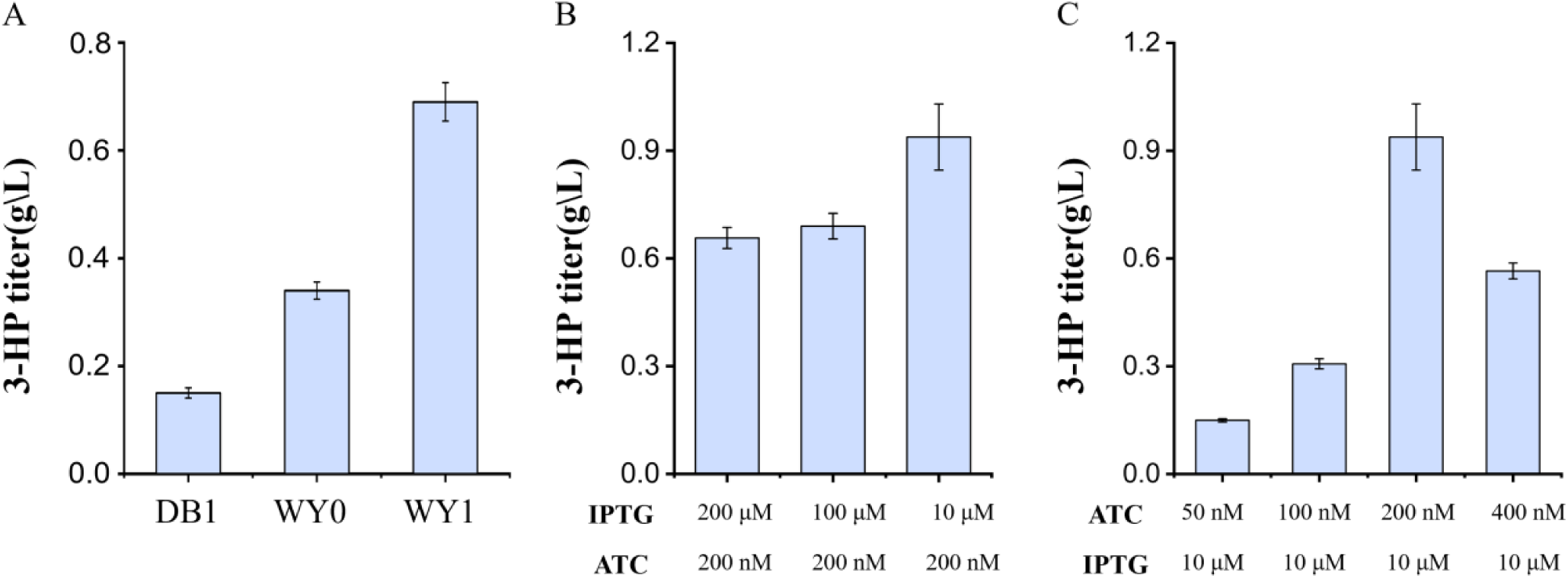
Effect of the induction strength on 3-HP production by strain WY1. (A) 3-HP titer of strains WY0, WY1 and DB1; (B) 3-HP titer of strain WY1 in response to IPTG concentration (10, 100, 200 μM); (C) 3-HP titer of strain WY1 in response to Tc concentration (50, 100, 200, 400 nM).

To optimize the culture conditions, we subsequently tried the different concentrations of inducers, and the 3-HP titer of strain WY1 varied with the different conc. of IPTG and Tc, results shown in Fig. 3B and Fig. 3C. Comparing the treatment groups, we found that strain WY1 obtained the highest 3-HP concentration of approximately 0.94 g/L when IPTG 10 μmol/L and Tc 200 nmol/L were used.

### 3.2 Increasing acetyl-CoA precursor supply by decreasing by-product carbon flux

Acetyl-CoA is a critical component of microbial carbon metabolism in the intracellular metabolic network. It links metabolic processes such as the TCA cycle, glycolysis, glyoxylate shun, lactate and acetate pathways, and these products use acetyl-CoA as a precursor molecule. Acetyl-CoA is a critical component of microbial carbon metabolism in the intracellular metabolic network [33, 34]. Acetyl-CoA also plays a pivotal role in synthesis of 3-HP via the malonyl-CoA pathway. Consequently, there is a significant research imperative and potential for practical applications in enhancing the synthesis flux of intracellular acetyl-CoA. [35]. Two strategies were employed to augment the supply of acetyl-CoA in the 3-HP synthesis pathway. In addition to directly overexpressing *acc* to provide more malonyl-CoA for 3-HP synthesis, we also reduce by-product carbon flux by deleting *ldhA*/*poxB*/*pta* genes encoding key enzymes of the metabolic bypass lactate and acetate pathway, thereby increasing the intracellular acetyl-CoA supply.

One of the by-products of aerobic fermentation is acetate, which is produced by the overflow metabolism of *E. coli*. However, the accumulation of acetate inhibits cell development and limits the amount of recombinant protein, resulting in a loss of carbon supply. Two principal pathways are responsible for the formation of acetate in the metabolic pathway of *E. coli*. The phosphotransacetylase and acetate kinase (*pta*/*ackA*) pathway is dominant during the exponential phase, while the pyruvate oxidase (*poxB*) pathway is dominant during the stationary phase [36]. In addition, lactate formation depends on the D-lactate dehydrogenase encoded by the *ldhA* gene [37].

We investigated the impact of single-gene deletion on the 3-HP titer. Compared the characteristics of recombined strains (DB1, WY1, WY2, and WY5) regarding the accumulation of the metabolites 3HP, lactate, and acetate during glucose consumption in shake-flask experiments, with results presented in Fig. 4.

**Fig. 4.**
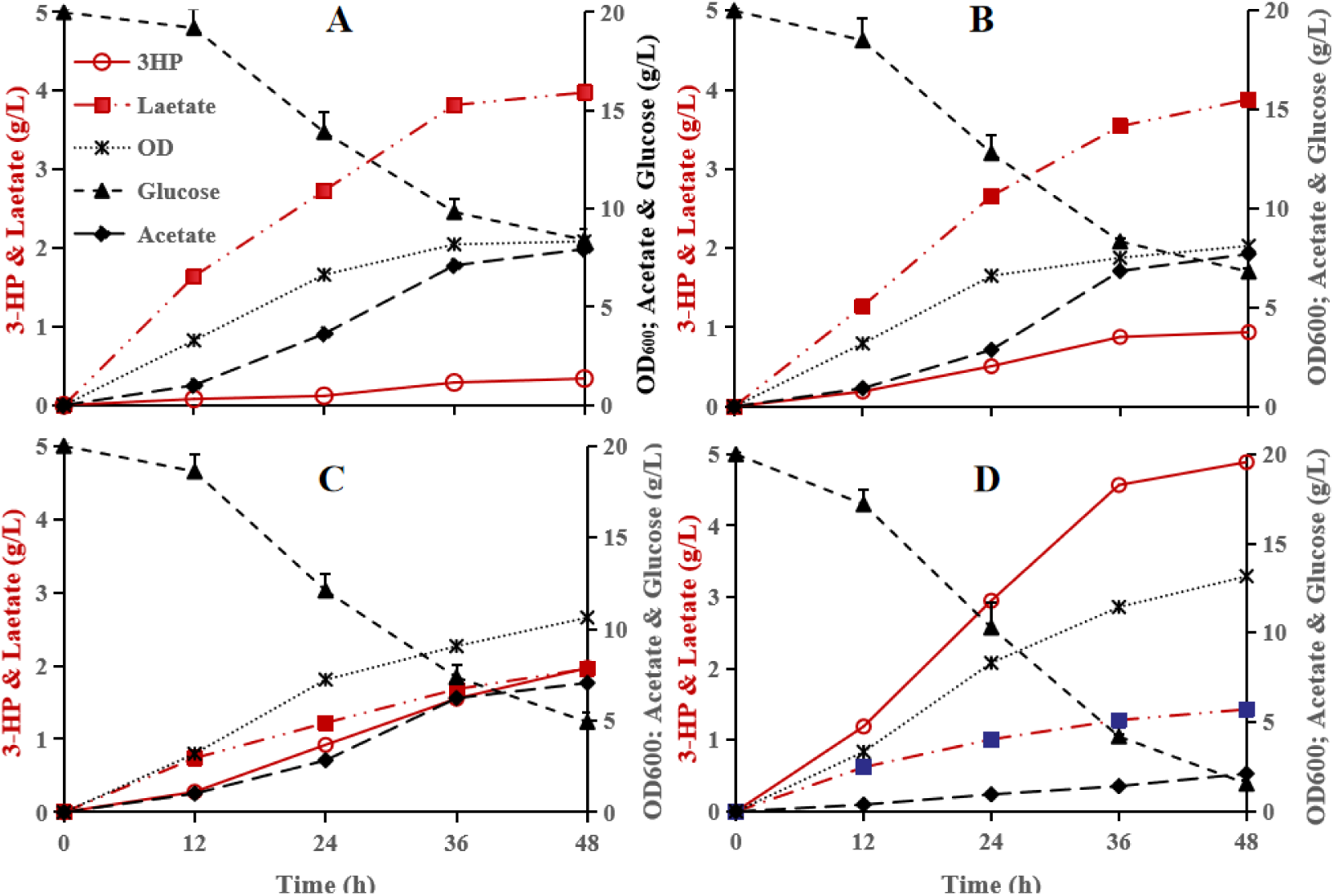
Time course of metabolite (3HP, lactate, acetate) accumulation, and glucose consumption and biomass changed by recombinant strains in shake flasks. A) DB1, strain was wild-type *E. coli* BL21 (DE3) with pA2s-mcr.B) WY1, strain BL21(DE3) with plasmid pE7a-acc and pA2s-mcr-N-C*, and 3 mutations were introduced into the segmented mcr.C) WY2, strain with deletion of genes *ldhA* based on strain WY1; D) WY5, strain Q1Z2 transformed with pA2s-mcr-N-C*, and Q1Z2 was BL21(DE3) Δ*ldhA* Δ*pta*::119-*accDABC* Δ*poxB*:: *CPA1*-*accDABCaccBC*.

The DB1 strain was wild-type *E. coli* BL21 (DE3) harbored pA2s-mcr. The DB1 produced 3HP, lactate, acetate was 0.34g/L, 3.98g/L, 7.95g/L, when fermented in modified M9 medium supplemented with 20 g/L glucose at 48 h, respectively ( Fig. 4A). The WY1 is based on the strain DB1 and harbored pE7a-acc, and introduced three mutation points in *mcr* segmented of strain DB1 according to Liu C (2016). Compared with the DB1, the 3HP titer increased 2.62 times more than WY1 (Fig. 4B). Compared to the lactate titer of 3.88 g/L produced by WY1, the strain WY2 based on the WY1 and deleted the *ldhA* gene, the lactate concentration decreased to 1.96 g/L in the WY2 fermentation broth, while the 3HP titer increased to 1.97 g/L from 0.94 g/L produced by WY1( Fig. 4C). And the 3HP titer of WY5 reached to 4.89 g/L when replaced with the original genes *ldhA*, *pta, poxB* on the genome with the genes *accDABC* and *accDABCaccBC*. The titer of 3HP increased by 4.2 times higher than strain WY1. This results demonstrated when reducing the branch from pyruvate to lactate or acetate carbon flow can significantly increase the 3HP yield. Furthermore, the key genes *accDABC,* especially the *accBC*, which encodes rate-limiting enzyme acetyl-CoA carboxylase, was integrated into the chromosome can obviously increase synthesis of downstream product 3HP. Compared with other strains, the WY5 harvested the highest 3-HP titer and cell density than DB1, WY1 and WY2,in the same condition(Tab. 2), suggesting that WY5 has a more carbon flow to synthesize 3-HP. These results indicated that knockout of the by-products synthesis pathway key genes *ldhA*, *poxB*, *pta* and expression of rate-limited genes 2 of *accDABC* and *accBC* in the genome successfully alleviated the rate-limiting step and enhanced the production of 3-HP. Our research results are similar to literature results (Zhang 2023.Song 2024), they decelerated glucose transport system to overcome acetate overflow and toxic intermediate 3-hydroxypropionaldehyde (3-HPA) accumulation, so as to alleviate growth retardation and improved the 3-HP biosynthesis.

**Tab. 2.** 3-HP production, residual glucose concentrationand cell density of strain DB1,WY1 and WY5.

| Strain name | Gene information | 3-HP titer (g/L) | Residual glucose (g/L) | 3-HP yield (g/g glucose) | OD<sub>600</sub> |
| --- | --- | --- | --- | --- | --- |
| DB1 | BL21(DE3)/pA2s-mcr | 0.34 | 8.42 | 0.029 | 8.34 |
| WY1 | DB2+pA2s-mcr-N-C* | 0.94 | 6.82 | 0.071 | 8.12 |
| WY2 | DB2+pA2s-mcr-N-C* Δ*ldhA* | 1.97 | 4.93 | 0.130 | 10.63 |
| WY5 | BL21(DE3)Δ*ldhA*Δ*pta*::119-*accDABC*Δ*poxB*::*CP A1-accDABCaccBC*+pA2s-mcr-N-C* | 4.89 | 1.56 | 0.261 | 13.17 |

\*Site-Directed Mutation on the carbon-terminal of *mcr* sequence

**Tab. 3.** Representative studies utilizing different pathways for the production of 3-HP biotechnology.

| Microorganisms | Pathway | Carbon Source | Engineering strategy | Titer (g/L) | Ref. |
| --- | --- | --- | --- | --- | --- |
| *L. reuteri*+*E. coli* | Coenzyme A-dependent pathway | Glycerol | Expression of *gabD4* and *pduQ* | 125.93 | [6] |
| *K. pneumoniae* | Coenzyme A-independent pathway | Glycerol | Expression of *puuC* by using tandem repeat tac promoter | 102.6 | [16] |
| *E. coli* | β-alanine pathway | Glucose | Expression of *pa0132*, *ydfG*, *panD*, *aspA* and *ppc*; deletion of *fumAC*, *fumB* and *iclR* | 31.1 | [40] |
| *E. coli* | Malonyl-CoA pathway | Glucose | Modulation of MCR activity | 40.6 | [21] |
| *E. coli* | Malonyl-CoA pathway | Glucose | Expression of *accDABC* and *mcr*; deletion of *ldhA*, *poxB* and *pta*; Biosensor-based ALE | 48.8 | This work |

### 3.3 Increased production of 3-HP via adaptive laboratory evolution using biosensor

It was previously demonstrated that the PpHpdR/PhpdH gene expression system, which is induced by 3-HP, exists in *P. denitrificans* [39]. In this system, 3-HP induces LysR-type transcriptional regulators to activate the expression of catabolic genes involved in the degradation of 3-HP. Based on this system, we developed a selection plasmid capable of recognizing 3-HP as a synthetic biosensor (Fig. 5 A, B, C, D). We amplified the gene encoding the red fluorescence protein (mCherry), the gene encoding chloramphenicol acetyltransferase (CmR), and the PpHpdR/PhpdH system from the corresponding plasmid templates. These genes were then inserted into the plasmid pYB01, with mCherry and CmR positioned downstream of the PpHpdR/PhpdH system.

**Fig. 5.**
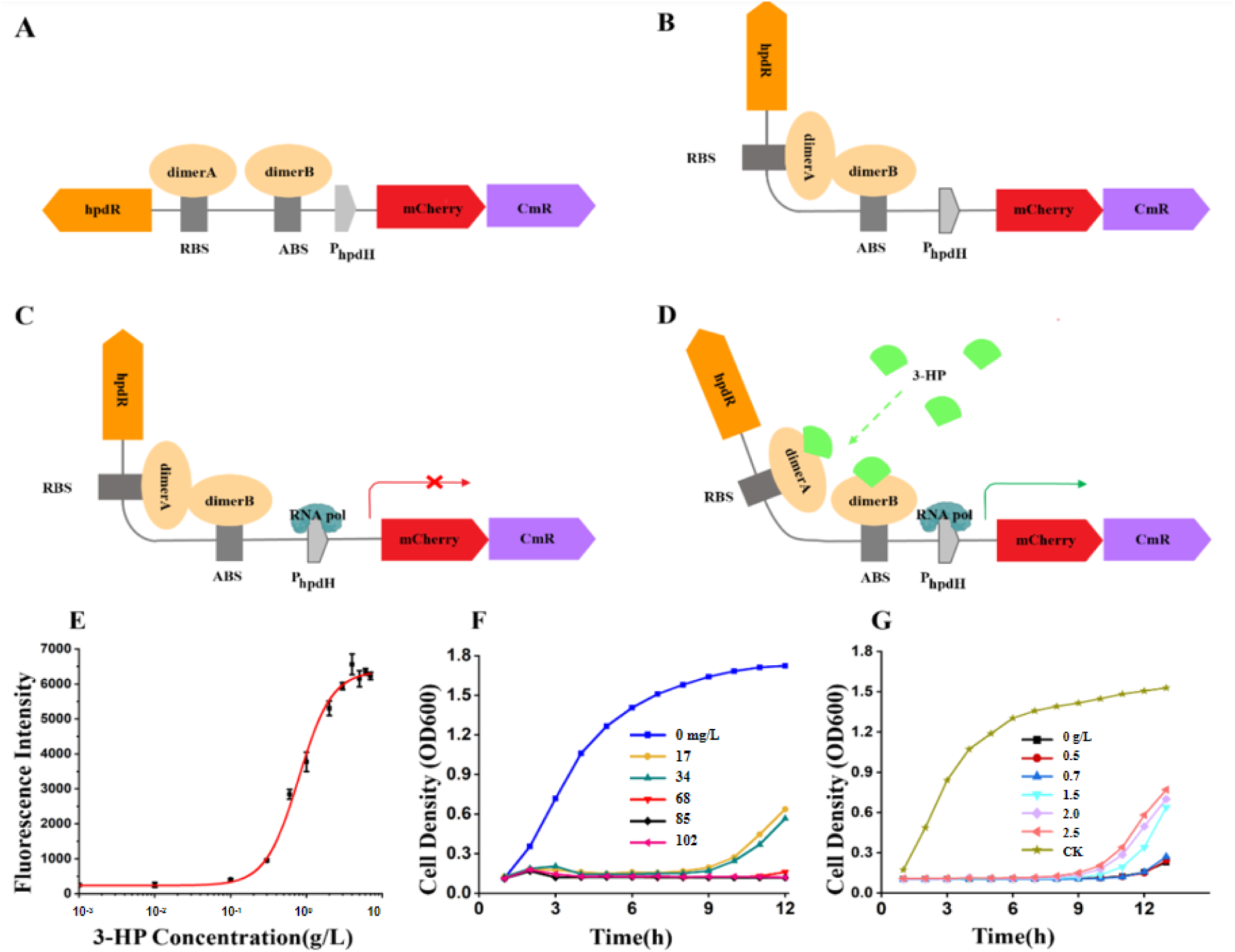
Schematic diagram of the working principle of the biosensor and validation of its effectiveness. (A) Dimers A and B produced by hpdR are combined on Regulatory binding site (RBS) and Activation Binding Site (ABS); (B) The interaction between the two dimers forms a tetramer, leading to DNA bending; (C) RNA polymerase binds to the promoter. Transcription cannot be activated because of the abnormal conformation of DNA; (D) As 3-HP exists, 3-HP binds to hpdR tetramer, protein conformation changes, DNA bending is alleviated, and transcription is activated; (E) The dose-response curve of 3-HP-induced expression *Pp*HpdR/P_hpdH_ in Q1Z2; The figure shows the relationship between the concentration of the inducer and fluorescence output after 3-HP was after 3-HP was added for about 12 hours. The inducer levels were 0.0001, 0.001, 0.01, 0.1, 0.2, 0.3, 0.4, 0.5,0.6 and 0.7 g/L, respectively. The error bars indicate the standard deviations of measurements from three independent cultures; (F) The cell density of strain CS2 at concentrations of chloramphenicol (0, 17, 34, 68, 85, 102 mg/L) when no 3-HP in medium; (G) The cell density of strain CS2 when adding concentrations of 3-HP (0.5, 0.7, 1.5, 2.0, 2.5 g/L), and the starting concentration of chloramphenicol in the medium is 17 mg/L, and the control group (CK) receives no additional chloramphenicol or 3-HP.

Therefore, we engineered the biosensor plasmid pYB-0055-mCherry-CmR. The successful synthesis of the plasmid was confirmed using colony PCR and single enzyme digestion (see supplementary Fig. S4). This biosensor induces CmR expression through 3-HP, allowing only those cells that efficiently convert glucose to 3-HP to survive under the selection pressure of chloramphenicol. As a result, the cells exhibiting advantageous mutations for 3-HP overproduction will progressively prevail under this selective pressure.

After the biosensor plasmid was successfully constructed, it was transferred into Q1Z2 to create a recombinant strain called CS2. To verify the efficacy of the constructed biosensor, we measured the 3-HP dose-fluorescence intensity curve of strain CS2. The results showed that the fluorescence intensity of strain CS2 increased gradually when the concentration of 3-HP in the culture medium increased from 0 to 7 g/L and reached the maximum when the concentration of 3-HP was 6 g/L (Fig. 5E). We evaluated the growth of the CS2 under different chloramphenicol concentrations (Fig. 5F). The results showed that the 17 mg/L chloramphenicol have reduced growth rate obviously, whereas 68 mg/L have severely inhibit cell growth of the strain CS2 to a large extent. From these date, we demonstrated that this reduction in the 3-HP-driven biosensor strain can be recovered by adding exogenous 3-HP (Fig. 5G). The above results suggest that in the presence of 3-HP, expression of *CmR* downstream of the PpHpdR/PhpdH system is activated, allowing strains possessing this plasmid to survive in chloramphenicol-containing media. These results indicated that this ALE biosensor can be used to improve the production of 3-HP.

To initiate ALE, the biosensor plasmid was introduced into the WY5 strain to generate the starting parental strain. An initial selection pressure of 34 mg/L chloramphenicol was applied, a concentration that significantly hindered the growth of the biosensor strain responsive to 3-HP. As shown in Fig. 6., in the first stage (cycles 1-11), the strain tolerated 34 mg/L chloramphenicol, reaching an OD_600_ of 1.05 after 12 h. In the second stage (cycles 11-23), the chloramphenicol concentration increased to 68 mg/L, with OD_600_ reaching 1.24 by cycle 23. During the third stage (cycles 25-52) at 119 mg/L chloramphenicol, the adaptive evolution rate slowed, requiring about 2 times longer to achieve similar OD_600_ as in the second phase, and reaching OD_600_ of 1.48 by cycle 52. From this process, a strain exhibiting enhanced tolerance and improved growth under the highest antibiotic concentration was isolated and designated as WY7.

**Fig. 6.**
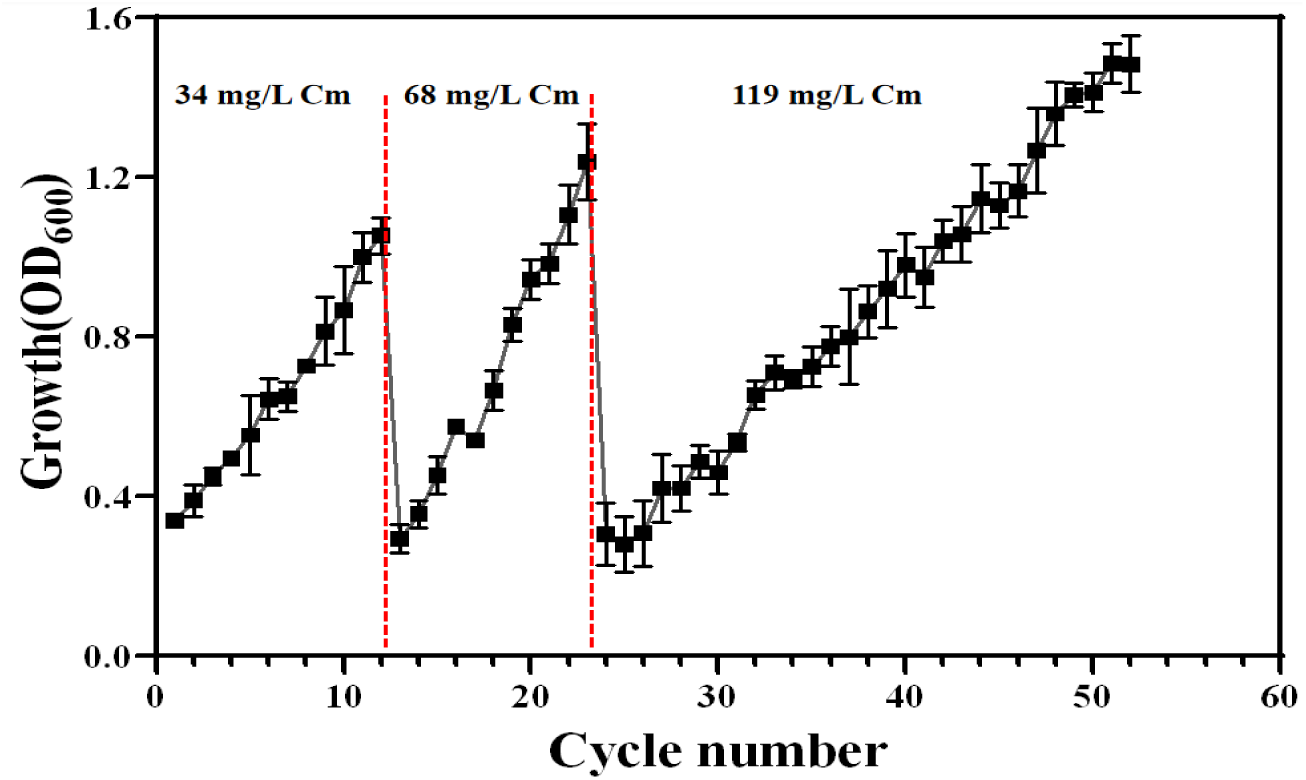
Growth profile of WY6 under 34-119 mg/L chloramphenicol during ALE from 1 to 52 cycles.

### 3.4. Evaluation of 3-HP production through shake flask and fed-batch fermentation

To verify synthesize 3HP ability of WY7, fermentation was conducted in 50 mL shake flask with 10 g/L modified M9 medium (20 g/L glucose, 2 g/L yeast extract) at 37℃ with shaking 200 rpm. After 2.5 h of induction, 40 mg/L biotin and 20 mM NaHCO_3_ were added to the medium to assist the expression of *acc.* WY7 fermentation broth were taken every 12 hours to measure the metabolite accumulation, glucose consumption and biomass dynamic changes in 48 hours, the results showed on Fig. 7A. WY7* accumulated 3HP, lactate and acetate were 6.17 g/L, 0.31 g/L and 1.15 g/L after 48 hours fermentation, respectively. Above results showed that the by-production lactate and acetate observably decreased than the wild type (Zhang YF, 2023).

**Fig. 7.**
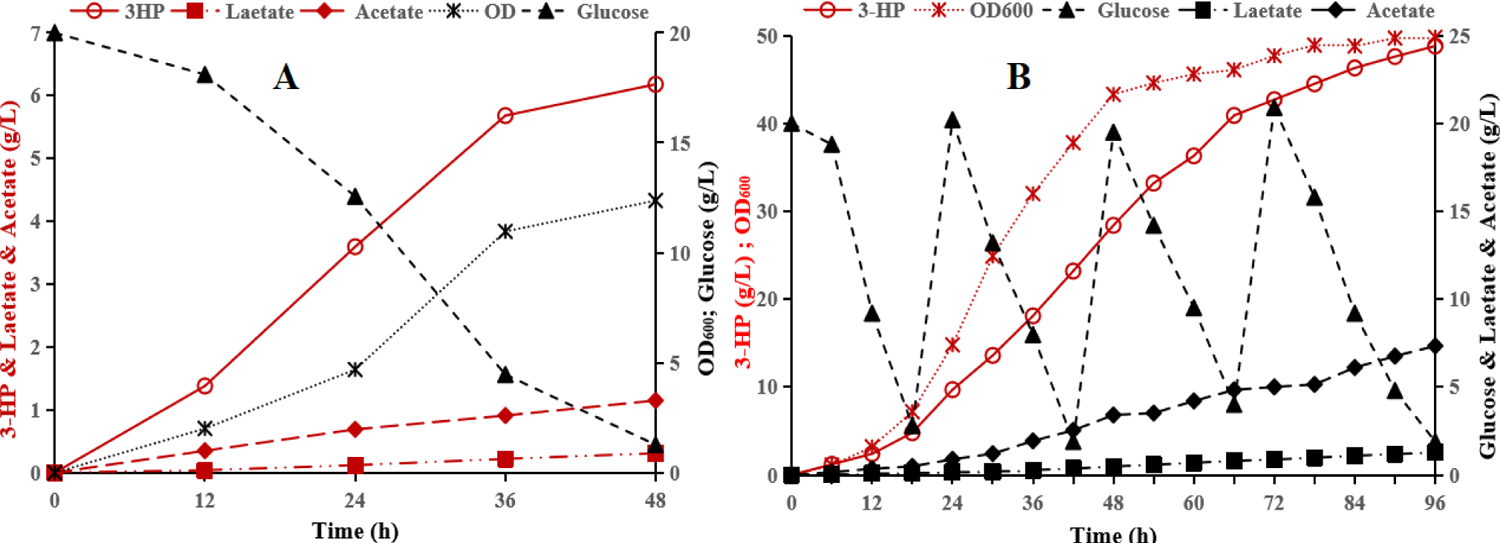
Time course of metabolite (3HP, lactate, acetate) accumulation, and glucose consumption and biomass changed by strain WY7* in shake flasks (A) and fed-batch(B) fermentation.

To fully verify the production potential of the strain WY7 and the production capacity of synthesize 3-HP in a large-scale process, we performed a 5 L fed-batch fermentation. Fed-batch fermentation was conducted to identify optimal synthesis conditions and maximum production based on an extended fermentation process condition. After performing a 96 hours fed-batch fermentation, a distinctly different metabolite profile was observed compared to short-duration batch fermentation. The final titer of 3-HP, lactate and acetate were 48.8 g/L, 1.28 g/L and 7.34 g/L, substantially (Fig. 7B). And the titer of 3HP increased 6.9 times higher than the short-time (48-h) batch fermentation. However, there were still 7.34 g/L of acetate in the fermentation broth, this result suggested that increasing the yield of 3HP can be achieved by knocking out reducing the competition for malonyl-CoA by inhibiting the fatty acids synthesis pathway, and enhanced the supply of NADPH and acetyl-CoA through cofactor engineering then accumulate 18.87 g/L 3-HP (Zhang WR, 2025).

## 4. Conclusions

3-HP has considerable economic value as a platform chemical. In the present study, two strategies were employed to genetically modify *E.coli* in order to facilitate the production of high concentrations of 3-HP via the malonyl-CoA pathway. First, we introduced the enzyme gene *mcr* from *Chloroflexus aurantiacus* into *E. coli* and overexpressed *acc*DABC from itself. For *mcr*, we divided it into two functional fragments to enhance its catalytic effect. In addition, the acetate synthesis pathway gene *pta/poxB* and the lactate synthesis pathway gene *ldhA* were knocked out to enhance precursor supply. On the basis of the above rational design, we introduced metabolite biosensors explicitly responsive to 3-HP into the engineered strains, coupled the 3-HP production to the growth of the strains, and promoted ALE production under the selective pressure of chloramphenicol. Finally, using glucose as a carbon source, our evolved strain WY7 accumulated the highest 3-HP titer of 48.8 g/L in a 5-liter fermenter, and the optimization of its culture conditions is still under intensive study.

## Declarations

**Ethics approval and consent to participate:** Not applicable

**Consent for publication:**Not applicable

## Availability of data and materials

The datasets used and/or analysed during the current study are available from the corresponding author on reasonable request.

All data generated or analysed during this study are included in this published article [and its supplementary information files]

## Competing interests

The authors declare that they have no competing interests

## Funding

The authors gratefully acknowledged the funding supported by the Natural Science Foundation of Heilongjiang Province [Grant Number ZHLJZR230150002]

## Authors’ contributions

Yanwei Wang and Chenmei Suo did the experiments, Youxin Cui and Jingyu Yang performed the literature search and wrote the manuscripts. Mutaz Mohammed Abdallah assisted manuscript polishing. Hongyi Yang and Pengchao Wang revised the manuscript, Lixin Li and and Changli Liu designed and supervised the project.

## Reference

1. Zabed HM, Akter S, Rupani PF, Akor J, Zhang Y, Zhao M, et al. Biocatalytic gateway to convert glycerol into 3-hydroxypropionic acid in waste-based biorefineries: Fundamentals, limitations, and potential research strategies. Biotechnol Adv. 2023 Jan-Feb;62:108075. 10.1016/j.biotechadv.2022.108075.

2. de Fouchécour F, Sánchez-Castañeda AK, Saulou-Bérion C, Spinnler HÉ. Process engineering for microbial production of 3-hydroxypropionic acid. Biotechnol Adv. 2018 Jul-Aug;36(4):1207–1222. 10.1016/j.biotechadv.2018.03.020.

3. Zhang WR, Zhang HJ, Wu H, High-Yield Biosynthesis of 3-Hydroxypropionic Acid from Acetate in Metabolically Engineered *Escherichia coli*. ACS Synthetic Biology. 2025, May, 14, (5) 16 2025,1654–1666. 10.1021/acssynbio.5c00030

4. Wang X, Cui Z, Sun X, Wang Z, Chen T. Production of 3-hydroxypropionic acid from renewable substrates by metabolically engineered microorganisms: a review. Molecules. 2023 Feb 16;28(4):1888. 10.3390/molecules28041888.

5. Chen T, Zhang Y, Yun J, Zhao M, Zhang C, Chen Z, Zabed HM, Sun W, Qi X. Bioproduction of 3-Hydroxypropionic Acid by Enhancing the Precursor Supply with a Hybrid Pathway and Cofactor Regeneration. ACS Synth Biol. 2024 Oct 18;13(10):3366–3377. doi: 10.1021/acssynbio.4c00427.

6. Zhang Y, Yun J, Zabed HM, Dou Y, Zhang G, Zhao M, et al. High-level co-production of 3-hydroxypropionic acid and 1,3-propanediol from glycerol: Metabolic engineering and process optimization. Bioresour Technol. 2023 Feb; 369: 128438. 10.1016/j.biortech.2022.128438.

7. Dishisha T, Pyo SH, Hatti-Kaul R. Bio-based 3-hydroxypropionic and acrylic acid production from biodiesel glycerol via integrated microbial and chemical catalysis. Microb Cell Fact. 2015 Dec 21;14:200. 10.1186/s12934-015-0388-0.

8. Lu ZY, Chen Z, Xiao WD, The biodegradable poly(3-hydroxypropionate) from acrylic acid via 3-hydroxypropionic acid: Studies on catalytic hydration and self-polycondensation. Chemical Engineering Journal, 2025 March Volume 507,160382 10.1016/j.cej.2025.160382

9. Çatıker, Efkan, Filik, Tahsin. Direct Synthesis of Hyperbranched Poly(acrylic acid-co-3-hydroxypropionate), International Journal of Polymer Science, 2015(1): 231059 10.1155/2015/231059

10. Fei P, Zhang WR, Wu H, Carbon-negative bio-production of short-chain carboxylic acids (SCCAs) from syngas via the sequential two-stage bioprocess by Moorella thermoacetica and metabolically engineered *Escherichia coli*. Bioresour Technol. 2025 Jan,Volume 416, 131714. 10.1016/j.biortech.2024.131714

11. Werpy, T., and Petersen, G. Top Value Added Chemicals from Biomass: Volume I-Results of Screening for Potential Candidates from Sugars and Synthesis Gas. United States: N. p., 2004.

12. Fina A, Brêda GC, Pérez-Trujillo M, Freire DMG, Almeida RV, Albiol J, Ferrer P. Benchmarking recombinant *Pichia pastoris* for 3-hydroxypropionic acid production from glycerol. Microb Biotechnol. 2021 Jul;14(4):1671–1682. doi: 10.1111/1751-7915.13833. Epub 2021 Jun 3.

13. Ju SB, Seo MJ, Yeom SJ. In Vitro One-Pot 3-Hydroxypropanal Production from Cheap C1 and C2 Compounds. Int J Mol Sci. 2022 Apr 3;23(7):3990. doi: 10.3390/ijms23073990.

14. Borodina I, Kildegaard KR, Jensen NB, Blicher TH, Maury J, Sherstyk S, et al. Establishing a synthetic pathway for high-level production of 3-hydroxypropionic acid in *Saccharomyces cerevisiae* via β-alanine. Metab Eng. 2015 Jan;27:57–64. 10.1016/j.ymben.2014.10.003.

15. Rathnasingh C, Raj SM, Lee Y, Catherine C, Ashok S, Park S. Production of 3-hydroxypropionic acid via malonyl-CoA pathway using recombinant *Escherichia coli* strains. J Biotechnol. 2012 Feb 20;157(4):633–40. 10.1016/j.jbiotec.2011.06.008.

16. Zhao P, Tian P. Biosynthesis pathways and strategies for improving 3-hydroxypropionic acid production in bacteria. World J Microbiol Biotechnol. 2021 Jun 15;37(7):117. 10.1007/s11274-021-03091-6.

17. Liu C, Ding Y, Xian M, Liu M, Liu H, Ma Q, et al. Malonyl-CoA pathway: a promising route for 3-hydroxypropionate biosynthesis. Crit Rev Biotechnol. 2017 Nov;37(7):933–941. 10.1080/07388551.2016.127209.

18. Liang B, Sun G, Zhang X, Nie Q, Zhao Y, Yang J. Recent advances, challenges and metabolic engineering strategies in the biosynthesis of 3-hydroxypropionic acid. Biotechnol Bioeng. 2022 Oct;119(10):2639–2668. 10.1002/bit.28170.

19. Liu D, Xiao Y, Evans BS, Zhang F. Negative feedback regulation of fatty acid production based on a malonyl-CoA sensor-actuator. ACS Synth Biol. 2015 Feb 20;4(2):132–40. 10.1021/sb400158w.

20. Wang S, Jin X, Jiang W, Wang Q, Qi Q, Liang Q. The expression modulation of the key enzyme acc for highly efficient 3-hydroxypropionic acid production. Front Microbiol. 2022 May 11;13:902848. 10.3389/fmicb.2022.902848.

21. Liu C, Ding Y, Zhang R, Liu H, Xian M, Zhao G. Functional balance between enzymes in malonyl-CoA pathway for 3-hydroxypropionate biosynthesis. Metab Eng. 2016 Mar;34:104–111. 10.1016/j.ymben.2016.01.001.

22. Tong T, Chen X, Hu G, Wang XL, Liu GQ, Liu L. Engineering microbial metabolic energy homeostasis for improved bioproduction. Biotechnol Adv. 2021 Dec;53:107841. 10.1016/j.biotechadv.2021.107841.

23. Marchetto F, Conde T, Kargul J, Adaptive laboratory evolution of extremophilic red microalga Cyanidioschyzon merolae under high nickel stress enhances lipid production and alleviates oxidative damage. Bioresour Technol. 2025 Oct, 434,132826, 10.1016/j.biortech.2025.132826

24. Sandberg TE, Salazar MJ, Weng LL, Palsson BO, Feist AM. The emergence of adaptive laboratory evolution as an efficient tool for biological discovery and industrial biotechnology. Metab Eng. 2019 Dec;56:1–16. 10.1016/j.ymben.2019.08.004.

25. Mo X, Zhao Y, Zhu L, Zhang C, Liu Z, Ma Z, Bao K, Yang S. Adaptively evolved *Methylorubrum extorquens* with enhanced formate tolerance and its application in 3-hydroxypropionic acid production. Appl Environ Microbiol. 2025 Aug 13:e0256024. doi: 10.1128/aem.02560-24.

26. Sun X, Peng Z, Li C, Zheng Y, Cheng Y, Zong J, et al. Combinatorial metabolic engineering and tolerance evolving of *Escherichia coli* for high production of 2’-fucosyllactose. Bioresour Technol. 2023 Mar;372:128667. 10.1016/j.biortech.2023.128667.

27. Kim K, Hou CY, Choe D, Kang M, Cho S, Sung BH, et al. Adaptive laboratory evolution of *Escherichia coli* W enhances gamma-aminobutyric acid production using glycerol as the carbon source. Metab Eng. 2022 Jan;69:59–72. 10.1016/j.ymben.2021.11.004.

28. Mundhada H, Seoane JM, Schneider K, Koza A, Christensen HB, Klein T, et al. Increased production of L-serine in *Escherichia coli* through Adaptive Laboratory Evolution. Metab Eng. 2017 Jan;39:141–150. 10.1016/j.ymben.2016.11.008.

29. Wang G, Li Q, Zhang Z, Yin X, Wang B, Yang X. Recent progress in adaptive laboratory evolution of industrial microorganisms. J Ind Microbiol Biotechnol. 2023 Feb 17;50(1):kuac023. doi: 10.1093/jimb/kuac023.

30. Lee TS, Krupa RA, Zhang F, Hajimorad M, Holtz WJ, Prasad N, et al. BglBrick vectors and datasheets: A synthetic biology platform for gene expression. J Biol Eng. 2011 Sep 20;5:12. 10.1186/1754-1611-5-12.

31. Gossen M, Freundlieb S, Bender G, Müller G, Hillen W, Bujard H. Transcriptional activation by tetracyclines in mammalian cells. Science. 1995 Jun 23;268(5218):1766–9. 10.1126/science.7792603.

32. Li C, Liu X, Li Y, Peng Q, Miao J, Liu X. The Tetracycline-Inducible/CRISPR-Cas9 System is an Efficient Tool for Studying Gene Function in *Phytophthora sojae*. Mol Plant Pathol. 2025 Jun;26(6):e70114. doi: 10.1111/mpp.70114.

33. Li X, Wang Y, Chen X, Eisentraut L, Zhan C, Nielsen J, Chen Y. Modular deregulation of central carbon metabolism for efficient xylose utilization in *Saccharomyces* cerevisiae. Nat Commun. 2025 May 16;16(1):4551. doi: 10.1038/s41467-025-59966-x.

34. Han X, Zhao Z, Wen Y, Chen Z. Enhancement of docosahexaenoic acid production by overexpression of ATP-citrate lyase and acetyl-CoA carboxylase in *Schizochytrium sp*. Biotechnol Biofuels. 2020 Jul 21;13:131. 10.1186/s13068-020-01767-z.

35. Takayama S, Ozaki A, Konishi R, Otomo C, Kishida M, Hirata Y, et al. Enhancing 3-hydroxypropionic acid production in combination with sugar supply engineering by cell surface-display and metabolic engineering of *Schizosaccharomyces pombe*. Microb Cell Fact. 2018 Nov 13;17(1):176. 10.1186/s12934-018-1025-5.

36. Ye T, Ding W, An Z, et al. Increased distribution of carbon metabolic flux during de novo cytidine biosynthesis via attenuation of the acetic acid metabolism pathway in *Escherichia coli*. Microb Cell Fact. 2025;24(1):36. doi:10.1186/s12934-025-02657-5

37. Cheng T, Liang X, Wang Y, Chen N, Feng D, Liang F, et al. Co-Production of Isoprene and Lactate by Engineered *Escherichia coli* in Microaerobic Conditions. Molecules. 2021 Nov 26;26(23):7173. 10.3390/molecules26237173.

38. Jiang Y, Chen B, Duan C, Sun B, Yang J, Yang S. Multigene editing in the Escherichia coli genome via the CRISPR-Cas9 system. Appl Environ Microbiol. 2015 Apr;81(7):2506–14. 10.1128/AEM.04023-14. Erratum in: Appl Environ Microbiol. 2016 May 31;82(12):3693. 10.1128/AEM.01181-16.

39. Zhou S, Ainala SK, Seol E, Nguyen TT, Park S. Inducible gene expression system by 3-hydroxypropionic acid. Biotechnol Biofuels. 2015 Oct 20;8:169. 10.1186/s13068-015-0353-5.

40. Song CW, Kim JW, Cho IJ, Lee SY. Metabolic Engineering of *Escherichia coli* for the Production of 3-Hydroxypropionic Acid and Malonic Acid through β-Alanine Route. ACS Synth Biol. 2016 Nov 18;5(11):1256–1263. 10.1021/acssynbio.6b00007.

41. Zhang YF, Yun JH, Zhang GY, Parvez A, Zhou L, Zabed H M, Li J, Qi XH, Efficient biosynthesis of 3-hydroxypropionic acid from glucose through multidimensional engineering of *Escherichia coli*. Bioresource Technology 389 (2023) 129822. 10.1016/j.biortech.2023.129822

42. Song C W, Kwon M, Park J M, Song H. Fermentative production of 3-hydroxypropionic acid by using metabolically engineered *Klebsiella pneumoniae* strains. Biochemical Engineering.212 (2024) 109516. 10.1016/j.bej.2024.109516

